# BRAT1 associates with INTS11/INTS9 heterodimer to regulate key neurodevelopmental genes

**DOI:** 10.1101/2023.08.10.552743

**Authors:** Sadat Dokaneheifard, Helena Gomes Dos Santos, Monica Guiselle Valencia, Harikumar Arigela, Ramin Shiekhattar

**Author notes:** Email addresses: S.D; H.G.D.S; M.G.V; H.A.

## Abstract

Integrator is a multi-subunits protein complex involved in regulation of gene expression. Several Integrator subunits have been found to be mutated in human neurodevelopmental disorders, suggesting a key role for the complex in the development of nervous system. *BRAT1* is similarly linked with neurodegenerative diseases and neurodevelopmental disorders such as rigidity and multifocal-seizure syndrome. Here, we show that INTS11 and INTS9 subunits of Integrator complex interact with BRAT1 and form a trimeric complex in human HEK293T cells as well as in pluripotent human embryonal carcinoma cell line (NT2). We find that *BRAT1* depletion disrupts the differentiation of NT2 cells into astrocytes and neural cells. Loss of *BRAT1* results in inability to activate many neuronal genes that are targets of REST, a neuronal silencer. We identified BRAT1 and INTS11 co-occupying the promoter region of these genes and pinpoint a role for BRAT1 in recruiting INTS11 to their promoters. Disease-causing mutations in *BRAT1* diminish its association with INTS11/INTS9, linking the manifestation of disease phenotypes with a defect in transcriptional activation of key neuronal genes by BRAT1/INTS11/INTS9 complex.

**Highlights:**

- Integrator subunits INTS9 and INTS11 tightly interact with BRAT1
- Depletion of *BRAT1* causes a dramatic delay in human neural differentiation
- BRAT1 and INTS11 module targets the promoters of neural marker genes and co-regulates their expression. The recruitment of INTS11 to these sites is BRAT1-dependent.
- Pathogenic E522K mutation in *BRAT1* disrupts its interaction with INTS11/INTS9 heterodimer.

## Introduction

*BRAT1* (BRCA1 Associated ATM Activator 1), also known as *BAAT1* (BRCA1-associated protein required for ATM activation 1), was initially described as a BRCA1-interaction partner and it is important for activation of ATM (Aglipay *et al*, 2006). It was reported that BRAT1 protein localizes to both nucleus and cytoplasm (Pourahmadiyan *et al*, 2021). Later studies suggested the involvement of BRAT1 in DNA damage response as well as MTOR signaling (Aglipay *et al*., 2006; Low *et al*, 2015; Ouchi & Ouchi, 2010; So & Ouchi, 2013, 2014). *BRAT1* depletion resulted in impaired cell cycle progression using synchronized mouse embryonic fibroblasts (MEF) under serum-starvation condition (So & Ouchi, 2013). Up-regulation of serum Anti-BRAT1 antibodies was indicated as a common molecular biomarker for gastrointestinal cancers and atherosclerosis (Hu *et al*, 2022). Although BRAT1 has been deemed un-druggable, recently the small molecule, Curcusone D, was shown to associate with BRAT1 and reduce cancer cells migration (Cui *et al*, 2021). *BRAT1* was found to be highly expressed in mouse brain (Ouchi & Ouchi, 2010). Consistent with functional importance of BRAT1 in the brain, germ line mutations in *BRAT1* are associated with rigidity and multifocal seizure syndrome, lethal neonatal and neurodevelopmental disorder with cerebellar atrophy and with or without seizure (Balasundaram *et al*, 2021; Mundy *et al*, 2016; Nuovo *et al*, 2022; Scheffer *et al*, 2019; Srivastava & Naidu, 2016; Srivastava *et al*, 2016). BRAT1 related genes are broadly found in Eukaryotes (source: eggnog v.6, ID: 5EM9B, 441 evolutionary related proteins)(Hernández-Plaza *et al*, 2023). While the protein domain composition is weakly conserved (source: Pfam (Mistry *et al*, 2021)), BRAT1 orthologs have a common theme of Leucine-rich repeats folding into helical structures (source: CATH classification 1.25.10.10 (Sillitoe *et al*, 2021)). Human BRAT1 has been described to contain a CIDE-N (cell death–inducing DFF45-like effector [CIDE] domain) at its N-terminus and two HEAT (Huntingtin, Elongation factor 3, A subunit of protein phosphatase 2A, and TOR1; aa 495-531, aa 544-576) repeats at its C-terminus(Ouchi & Ouchi, 2010).

Integrator is an evolutionarily conserved complex in metazoans containing at least 16 subunits(Baillat *et al*, 2005; Beckedorff *et al*, 2020; Dasilva *et al*, 2021; Pfleiderer & Galej, 2021) associated with the C-terminal domain (CTD) of RNA polymerase II (RNAPII)(Albrecht *et al*, 2018; Baillat *et al*., 2005; Beckedorff *et al*., 2020; Dasilva *et al*., 2021; Wu *et al*, 2017). Since its discovery, Integrator has been shown to be implicated in multiple pathways involving coding and noncoding RNA processing and transcriptional regulation(Kirstein *et al*, 2021). Recent studies have highlighted a role for Integrator in transcriptional initiation, pause release, elongation, as well as termination of protein-coding genes (Gardini *et al*, 2014; Skaar *et al*, 2015; Stadelmayer *et al*, 2014). Additionally, Integrator was shown to be involved in biogenesis of enhancer RNAs (eRNAs) (Lai *et al*, 2015) and transcriptional responsiveness to epidermal growth factor (EGF) signaling (Lai *et al*., 2015). Integrator was shown to play diverse roles in ciliogenesis (Jodoin *et al*, 2013), embryogenesis (Kapp *et al*, 2013), human lung function (Obeidat *et al*, 2013), adipose differentiation (Otani *et al*, 2013), and maturation of viral microRNAs (Cazalla *et al*, 2011; Xie *et al*, 2015). We recently pinpointed a function for Integrator in the maintenance of canonical mature microRNAs stability through facilitating the loading of AGO2 (Kirstein *et al*, 2023). Importantly, mutations in human Integrator complex have been linked to brain developmental syndrome and cerebellar atrophy (Mascibroda *et al*, 2020; Oegema *et al*, 2017).

The key catalytic subunit of Integrator, INTS11, forms a heterodimer with INTS9(Dominski *et al*, 2005) mediated by the C-terminal regions of each protein module(Wu *et al*., 2017). INTS4, a HEAT repeat-containing protein, was determined to be a direct interaction partner of the INTS9/INTS11 heterodimer forming the minimal Integrator cleavage module(Pfleiderer & Galej, 2021). INTS4 was suggested to play a ‘Symplekin-like’ scaffold making specific and conserved interaction with INTS9/INTS11 (Albrecht *et al*., 2018). We find that BRAT1 associates with INTS9/INTS11 heterodimer, and this complex does not contain INTS4 protein. Indeed, our molecular modeling suggests that BRAT1 folds into a helical “O” shape structure wrapping around the INTS9/INTS11 heterodimer which partially overlaps with the interface that binds INTS4. Therefore, we find BRAT1 and INTS4 form mutually exclusive complexes with INTS11/INTS9 heterodimer. Importantly, disease-causing mutations of BRAT1 display diminished interaction with INTS11/INTS9. We show that during neuronal differentiation of NT2 cells, BRAT1 occupies and regulates the expression of a set of genes critical for neurodevelopment. Taken together, our study reveals an association of BRAT1 with INTS11/INTS9 proteins to regulate a set of key neuronal genes involved in differentiation of NT2 cells into a neuronal fate.

## Results

### Integrator subunits INTS9 and INTS11 tightly interact with BRAT1

We discovered integrator as a complex that associates with the C-terminal domain (CTD) of RNA polymerase II (RNAPII)(Baillat *et al*., 2005; Kirstein *et al*., 2021). To fully define the Integrator complex subunit composition, we isolated Flag-INTS11 from HEK293T-derived stable cell line and subjected the affinity-purified fractions to mass spectrometry. Beyond the core components of integrator complex described previously (Baillat *et al*., 2005), we obtained sequences corresponding to BRAT1 protein (Fig. 1a and Sup. Table 1). Western blot analysis of affinity purified fractions confirmed the specific association of BRAT1 with components of Integrator complex isolated through affinity purification of INTS11 (Fig. 1b). We next developed a HEK293T-derived cell line stably expressing Flag-BRAT1 to rigorously define the composition of the BRAT1-containing complex(s) (Fig. 1c). Affinity purification of Flag-BRAT1 followed by western blot analyses and silver staining (Fig. 1c), identified the core catalytic subunits of Integrator complex, INTS11 and INTS9, as the key components of the BRAT1-containing complex. Beyond, INTS11 and INTS9, we did not find any other subunits of Integrator complex in association with BRAT1 (Fig. 1c). To define the molecular mass of the BRAT1/INTS11/INTS9 complex, we subjected the affinity purified BRAT1 preparation to Superose 6 gel filtration chromatography (Fig. 1d) followed by mass spectrometry (Fig. 1d and Sup. Table 2). BRAT1 protein eluted with INTS11 and INTS9 at fraction 34 which was confirmed by western blot analyses, silver staining and mass spectrometry (Fig. 1d and Sup. Table 2).

**Figure 1.**
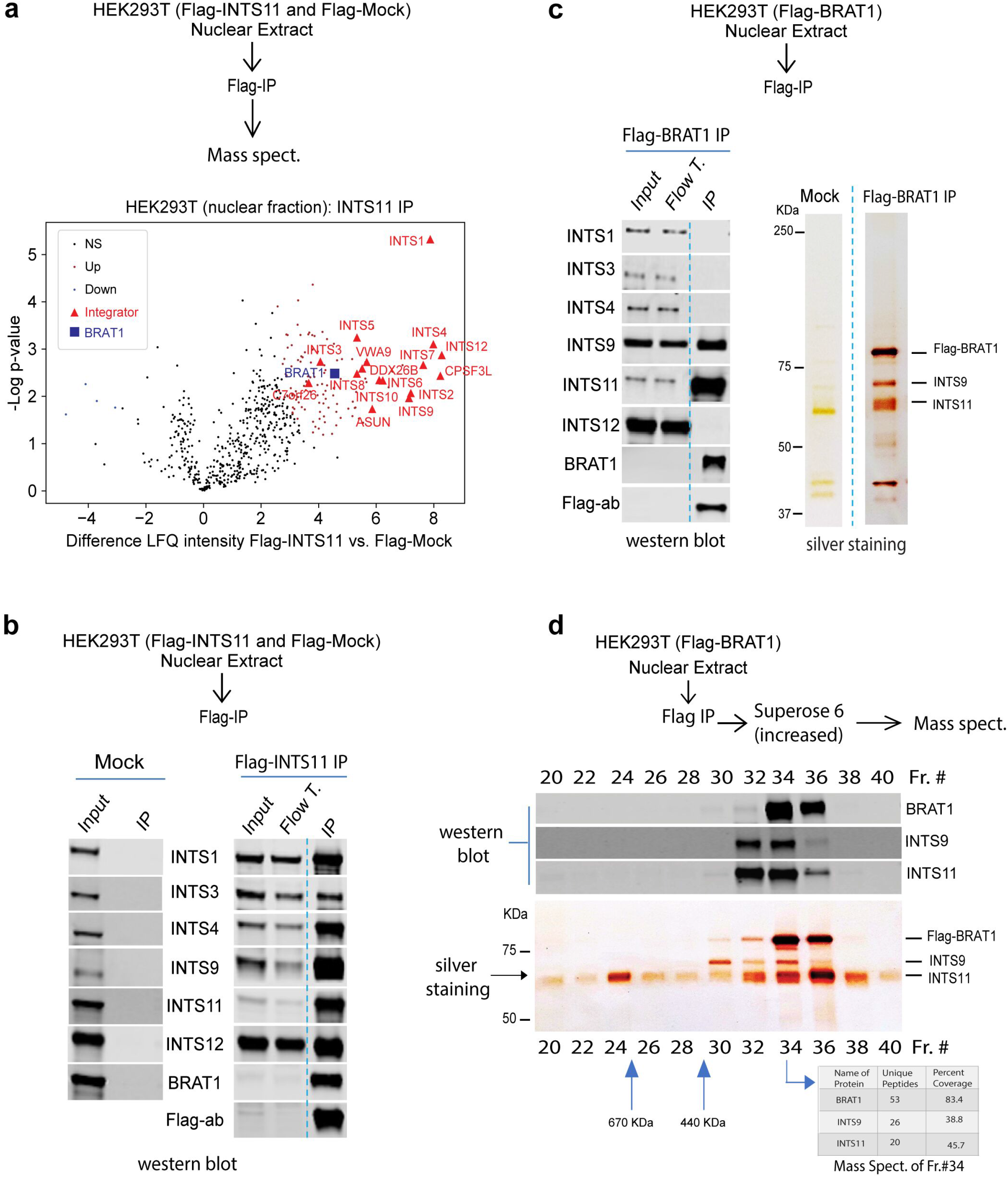
BRAT1 forms a stable interaction with INTS9/INTS11 heterodimer. a, Purification scheme and volcano plot of the protein enrichment in nuclear extract of HEK293T cells detected by mass spectrometric label-free quantification (LFQ) for Flag-INTS11 vs. mock (2 replicates per condition). Enriched proteins are shown as red circles and depleted proteins as blue circles, no significant protein interactors are shown as black circles. The enrichment levels of the full Integrator complex (red triangles) and BRAT1 (blue square) are highlighted (gene nomenclature: CPSF3L=INTS11, ASUN=INTS13, VWA9=INTS14). b, Western blot of affinity-purified Flag-INTS11 and Flag-mock. c, Western blot and silver staining of affinity-purified Flag-BRAT1 from nuclear lysate of HEK293T cells stably expressing Flag-BRAT1. Mock was used as control. d, Affinity-purified Flag-BRAT1 from nuclear extract of HEK293T cells stably expressing Flag-BRAT1 was fractionated on Superose 6 increased gel filtration and then visualized by western blot (top side) and silver staining (bottom side). Fraction numbers are indicated on top and bottom of the blot. Mass spectrometry on fraction 34 of nuclear extract of HEK293T cells stably expressing Flag-BRAT1 fractionated on Superose 6 increased gel filtration is shown by the table. Result confirmed the co-elution of BRAT1 with INTS11/INTS9 heterodimer in fraction 34. Number of unique peptides and their percent coverage for BRAT1/INTS11/INTS9 interactors are provided in the illustrated table.

**Table 1.**
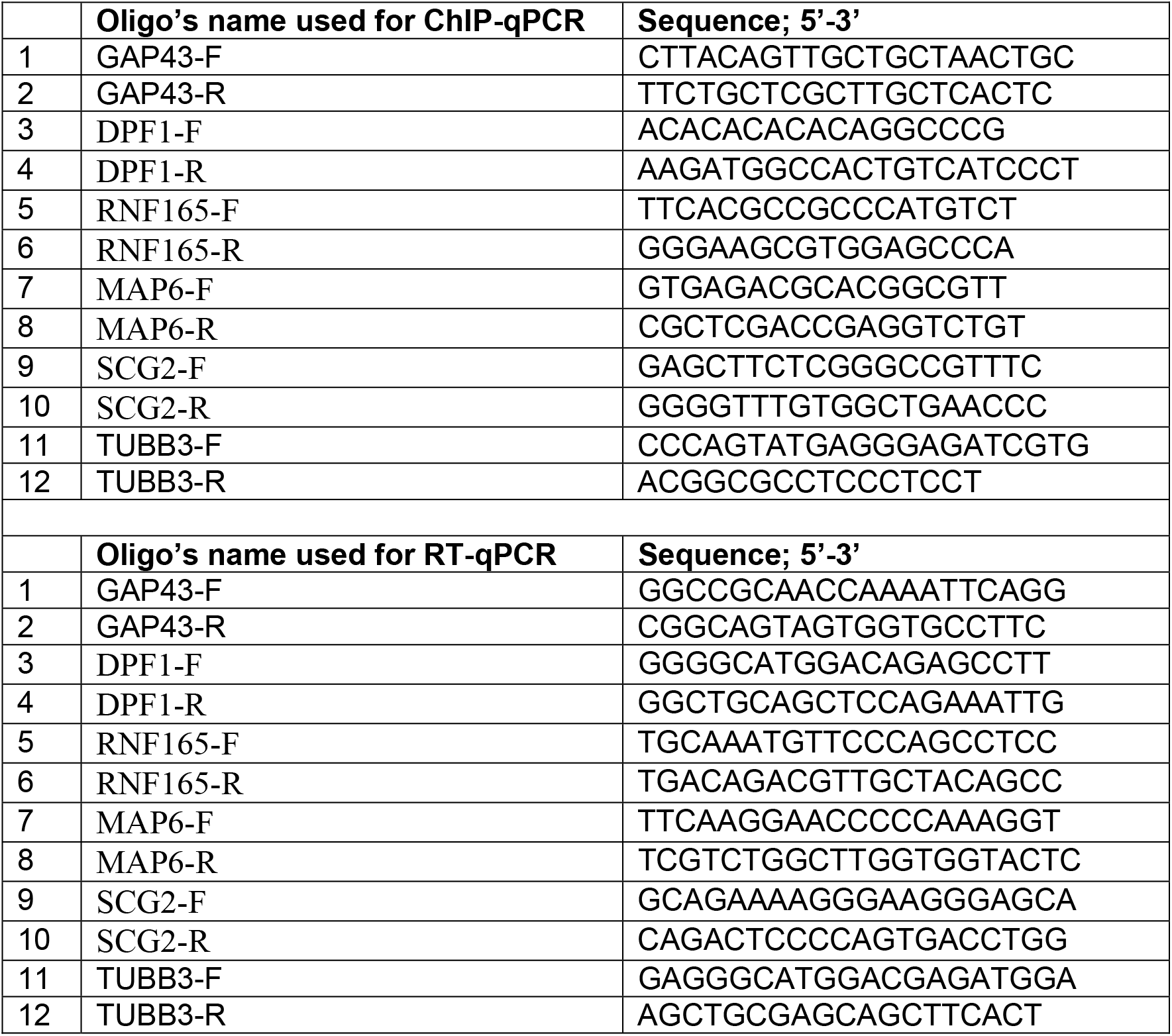
List of oligos used in this study.

### BRAT1 interacts with INTS11/INTS9 in NT2 cells, and it is required for the differentiation of NT2 cells into neurons and astrocytes

NTERA-2 (also known as NT2) cells are clonally derived pluripotent human embryonal carcinoma cell line that can be differentiated into a neuronal lineage following exposure to all-trans retinoic acid (ATRA) (Abolpour Mofrad *et al*, 2016; Andrews, 1984). Since, BRAT1 is associated with neurodevelopmental disorders (Balasundaram *et al*., 2021; Mundy *et al*., 2016; Nuovo *et al*., 2022; Scheffer *et al*., 2019; Srivastava & Naidu, 2016; Srivastava *et al*., 2016), we sought to determine the interaction between BRAT1 and INTS11/NTS9 in NT2 cells. We performed endogenous immunoprecipitation (IP) followed by western blot analysis (Fig. 2a) in NT2 cells. BRAT1 antibody immunoprecipitated INTS11 and INTS9, confirming our results from HEK293T cells. Additionally, immunoprecipitated eluate using INTS11 and INTS9 antibodies contained not only BRAT1 but also INTS4, reflecting the association of INTS11/INTS9 with two distinct complexes of Integrator and a second smaller complex containing BRAT1 (Fig. 2a).

**Fig. 2.**
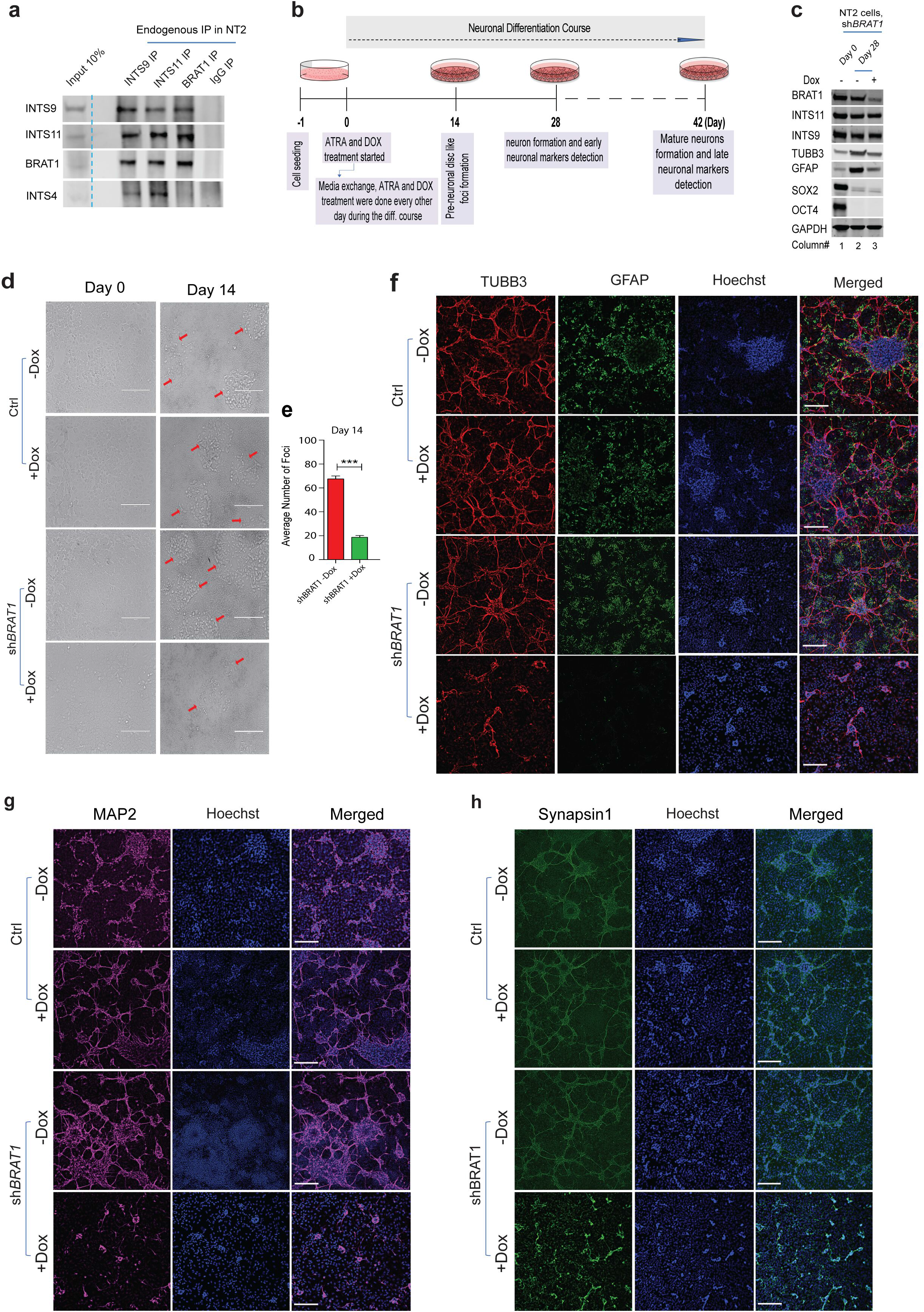
BRAT1 associates with INTS9/INTS11 heterodimer in NT2 cells and it is required for differentiation of NT2 cells into neurons and astrocytes. a, Immunoprecipitation of endogenous BRAT1, INTS11 and INTS9 in NT2 cells. The immunoblots of antibodies against INTS11 and INTS9 co-immunoprecipitated with BRAT1 and INTS4. Reciprocally, immunoblot antibody against BRAT1 co-immunoprecipitated only with INTS9 and INTS11. IgG was used as the negative control. Data shows the presence of the trimeric BRAT1/INTS11/INTS9 complex in NT2 cells. b, Diagram of the ATRA-treated based protocol used for the differentiation of NT2 cells. c, Western blot of BRAT1, INTS9 and INTS11 in parallel with stemness, neural and astrocyte markers 28 days post-differentiation of NT2 cells in BRAT1-depleted cells compared to control cells. Data shows that TUBB3 and GFAP protein levels decreased following BRAT1 depletion (lane 3) compared to non-depleted BRAT1 cells (lane 2) 28 days following ATRA treatment. d, Cell plate images showing the changes in cluster formation (bright and disc-like structures, highlighted by the red arrows) of NT2 cells at 14-day post-differentiation for BRAT1-depleted cells compared to control cells. Cells were imaged BF (bright field) with a EVOS XL Core Imaging System (Life Technologies) by a 20x objective. Final resolution figures were processed by ImageJ Fiji online tool (https://ij.imjoy.io/). Scale bar is 200 um. e, Bar plot of the average number of cell clusters (foci or bright and disc-like structures) per area at day 14, with Nikon TMS by a 4x objective. Data shows the average number of the clusters per area are significantly reduced in BRAT1-depleted cells compared to non-depleted cells 28 days post-differentiation (***p<0.001). f, Representative immunofluorescence images of TUBB3 (red, Biolegend #801201), GFAP (green, Invitrogen, #13-0300), Hoechst (blue), and merged images in NT2 cells 28 days post-differentiation. g, Representative immunofluorescence images of MAP2 (magenta, Millipore #AB5622), Hoechst (blue), and merged images in NT2 cells 28 days post-differentiation. h, Representative immunofluorescence images of Synapsin I (green, Synaptic Systems #106 011C3), Hoechst (blue), and merged images in NT2 cells 42 days post-differentiation. The scale bar for the microscopy images in panels f-h is 200 μm.

NT2 cells exposed to ATRA for 14 days develop into a pre-neuronal disc like foci (Fig. 2b). A more prolonged (28 days) exposure of NT2 cells to ATRA results in the appearance of neuronal and glial phenotypes (Fig. 2b). To assess the role of BRAT1 in neurogenesis, we depleted BRAT1 using an inducible short hairpin RNA (shRNA) in NT2 cells while treating the cells with ATRA to determine their differentiation into a neuronal and a glial cell fate (Fig. 2b). Importantly, depletion of BRAT1 in NT2 cells did not result in any changes in the protein Ievels of Integrator subunits as measured by western blot analyses (Fig. 2c and Sup. Fig. 2). Notably, we detected an increase in a TUBB3, a neural marker (Abolpour Mofrad *et al*., 2016; Conti *et al*, 2005; Shao *et al*, 2017) and GFAP, an astrocyte (glial) marker (Abolpour Mofrad *et al*., 2016; Jurga *et al*, 2021), following 28 days of ATRA treatment (Fig. 2c, lane 2). Additionally, we observed a reduction in OCT4 and SOX2, as the key stem cell factors, reflective of a loss of stemness and a gain of neuronal and glial phenotypes (Fig. 2c, lane 2). Critically, depletion of BRAT1 during the differentiation protocol led to a decreased expression of both TUBB3 and GFAP (Fig. 2c, lane 3). Consistent with the loss of TUBB3 and GFAP, depletion of BRAT1 decreased pre-neuronal foci formation following 14 days of ATRA treatment (Fig. 2d and e). Indeed, depletion of BRAT1 abrogated the establishment of neuronal and astrocyte phenotypes as measured by TUBB3, GFAP or MAP2 (Abolpour Mofrad *et al*., 2016; Conti *et al*., 2005) expression by day 28 or the late neuronal marker, Synapsin1 (Abolpour Mofrad *et al*., 2016), after 42 days of ATRA treatment (Fig. 2f-h). Taken together, these results point to a role for BRAT1 in neuronal and astrocytes differentiation in human NT2 cells.

### BRAT1 depletion leads to a defect in the expression of critical genes for neuronal function

To determine BRAT1-mediated changes in gene expression during the ATRA-induced differentiation of NT2 cells, we depleted BRAT1 and analyzed the transcriptome of NT2 cells. ATRA treatment in the control cells resulted in the differential expression of 11570 genes following 28 days where 5687 genes (49%) were down-regulated and a similar number of 5883 genes (51%) were up-regulated (1.5 fold-change and false discovery rate FDR<0.05) (Sup. Fig. 3a). Next, we depleted BRAT1 during the 28 days of differentiation and assessed the differential gene expression in NT2 cells undergoing neurogenesis. Remarkably, the loss of BRAT1 culminated in the decreased expression of a relatively small set of genes (250) (Fig.3a and b, Sup. Table 3). Critically, the prominent number of down-regulated genes play key roles in neuronal function including synaptic transmission and axonal guidance (Fig. 3c, Sup. Table 4). In contrast, differentially up-regulated genes (126) (Fig.3a and b, Sup. Table 3) control extracellular matrix organization and proliferation functions distinct from neuronal phenotype (Sup. Fig. 3b). These gene expression changes were validated using RT-qPCR (Fig. 3d-h). Taken together, our data suggest that BRAT1 is important for the expression of the genes involved in neuronal functions.

**Fig. 3.**
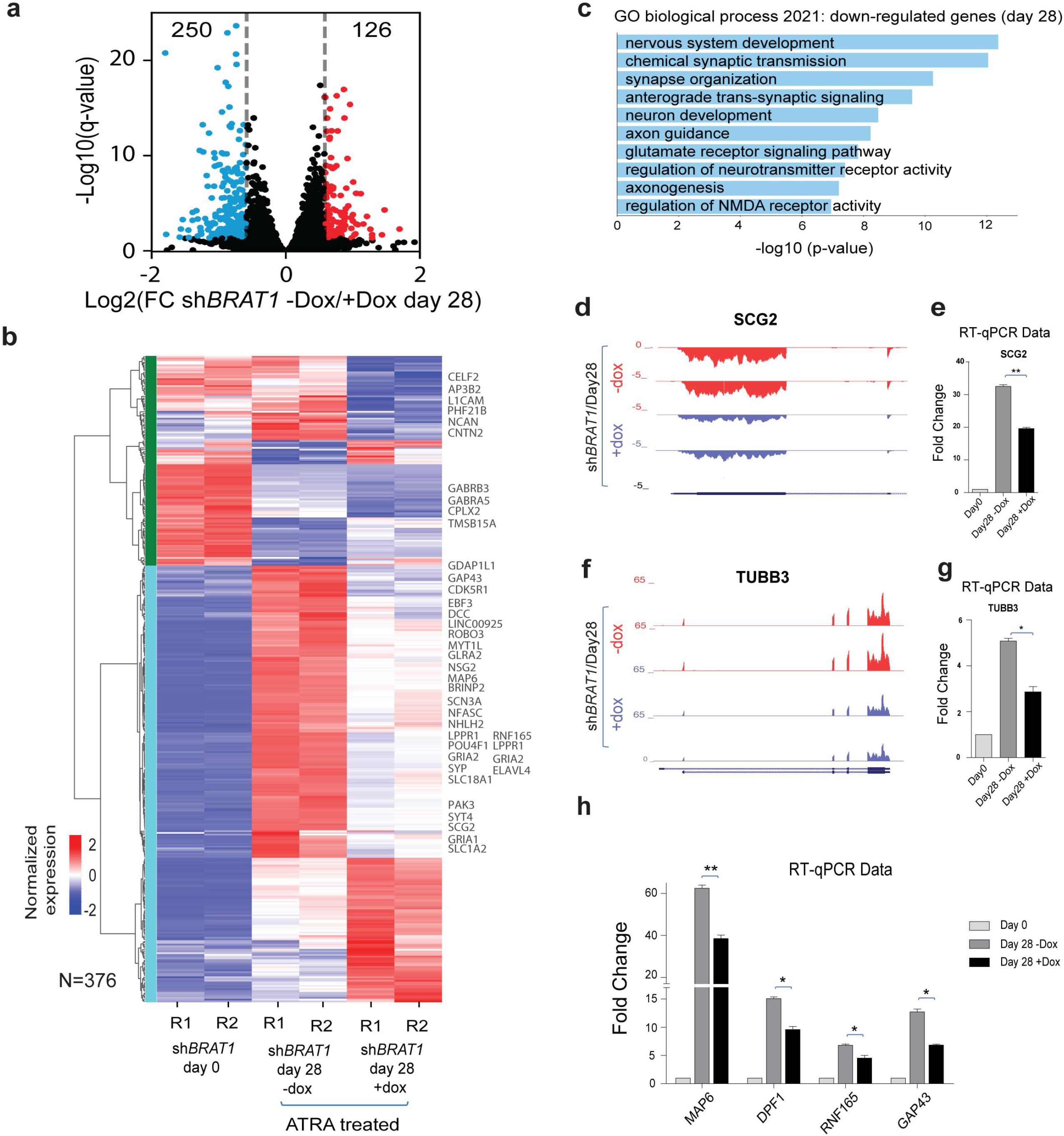
BRAT1 mediates the expression of a critical neurodevelopmental genes. a, Volcano plot of BRAT1-depleted cells compared to the cells expressing normal level of BRAT1 28 days post-ATRA treatment of NT2 cells, with down-regulated genes in blue, up-regulated genes in red and non-significant gene changes in black (significance cutoffs: 1.5-fold change and FDR<0.05). b, Heat map of the 376 genes differentially expressed from panel a. showing the row normalized gene expression levels (z-score) in control and BRAT1-depleted cells at day 0 (pre-treatment) and day 28 post-differentiation with ATRA treatment. Left color bar represent the clusters obtained by unsupervised hierarchical clustering (method = ward, metric = Euclidean distance). Genes listed on the right side are involved in neural development. c, Gene enrichment of the downregulated genes against Gene Ontology (GO) biological process 28 days after BRAT1 depletion in differentiated NT2 cells compared to the cells expressing normal level of BRAT1. Of the top ten GO biological processes most are categories associated to the nervous system including nervous system development, chemical synaptic transmission, and synapse organization. d, Genome browser tracks of the normalized RNA-seq data at SCG2 gene locus (28-days post-ATRA treatment, control cells expressing normal level of BRAT1 in red (-Dox), and BRAT1-depleted cells in blue (+Dox), 2 replicates each). e, Bar plot of the fold change in SCG2 gene expression of each sample compared to control by RT-qPCR (2 replicates). Expression was significantly decreased after BRAT1 depletion (+Dox) compared to the cells expressing normal level of BRAT1 (-Dox) at day 28 of ATRA treatment. f, Genome browser tracks of the normalized RNA-seq data for TUBB3 gene locus (28-days post-ATRA treatment, control cells expressing normal level of BRAT1 in red (-Dox), and BRAT1-depleted cells in blue (+Dox), 2 replicates each). g, Bar plot of the fold change in TUBB3 gene expression of each sample compared to control by RT-qPCR (2 replicates). Expression was significantly decreased after BRAT1 depletion (+Dox) compared to the cells expressing normal level of BRAT1 (-Dox) at day 28 of ATRA treatment. h, RT-qPCR data for showing additional examples of neural marker genes whose expressions are decreased 28 days post-differentiation in BRAT1-depleted cells (+Dox) compared to cells expressing normal level of BRAT1 (-Dox). For RT-qPCR experiments, data are expressed as ± SEM (2 repeats). Statistical significance was determined by unpaired t-test (*p < 0.05, **p<0.01).

### BRAT1 regulates INTS11 residence at critical neuronal differentiation genes

To determine the dynamic changes for BRAT1 and INTS11 proteins during the differentiation of NT2 cells into the neural lineage, we performed chromatin immunoprecipitation quantitative polymerase chain reaction (ChIP-qPCR) following ATRA-mediated differentiation (Fig. 4a). While ChIP-qPCR indicated the occupancy of BRAT1 and INTS11 at the promoter region of neural genes prior to stimulation with ATRA, we found a significant increase in INTS11 and BRAT1 residence at genes induced by ATRA following the differentiation protocol (Fig. 4a). We next asked whether BRAT1 regulates the occupancy of INTS11 at the promoter of BRAT1-responsive genes. ChIP-qPCR was performed using specific antibodies against BRAT1 and INTS11 in NT2 cells 3 days after BRAT1 depletion (Fig. 4b). Importantly, depletion of BRAT1 led to a significant reduction of INTS11 occupancy (Fig. 4b). Taken together, these results demonstrate that BRAT1 and INTS11 are concomitantly recruited during neural differentiation of NT2 cells to control the expression of the neural genes, and BRAT1 regulates the residence of INTS11 at the promoter of BRAT1-responsive genes.

**Fig. 4.**
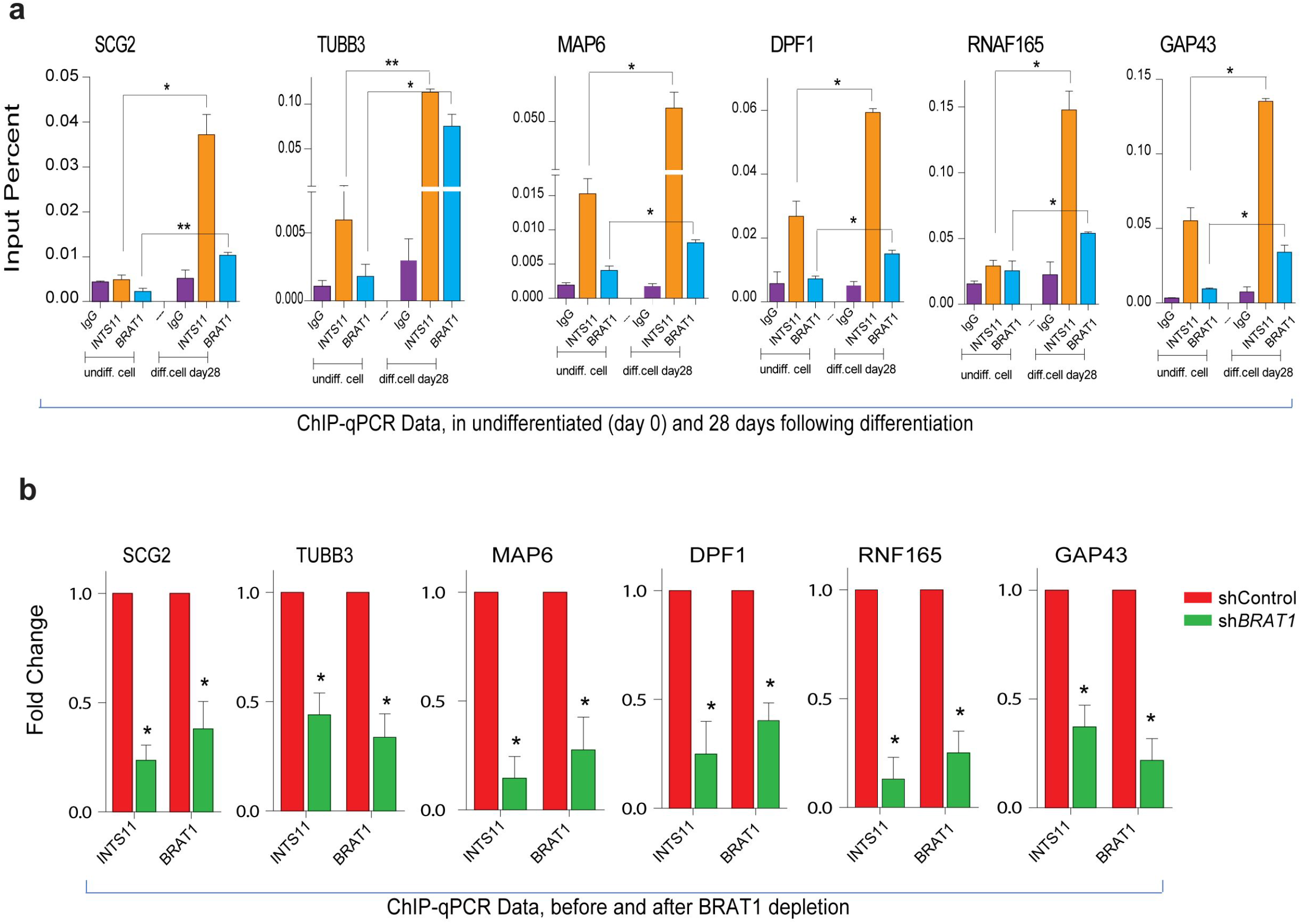
BRAT1 and INTS11 co-occupy the promoter of neuronal genes. a, Bar plots of INTS11 and BRAT1 ChIP-qPCR data at day 0 (undifferentiated cells) and at day 28 post-differentiation of NT2 cells showing significant increase of BRAT1 and INTS11 co-occupancy at the promoter region of neuronal genes activated following 28 days of ATRA treatment (normal differentiation). Each data in differentiated condition has been compared with its equivalent in undifferentiated condition. b, Bar plots of INTS11 and BRAT1 ChIP-qPCR data reveals the reduced enrichment of BRAT1 and INTS11 at the promoter region of neural marker genes following BRAT1 depletion (3 days after adding Dox without ATRA treatment) in sh*BRAT1* NT2 cells compared to shControl cells. For ChIP-qPCR, data are expressed as ± SEM fold change over shControl cells (2 repeats). Statistical significance was determined by unpaired t-test (*p < 0.05, **p<0.01).

### Pathogenic E522K mutation in BRAT1 disrupts its interaction with INTS11/INTS9 heterodimer

Recent reports link mutations in human Integrator subunits to neurodevelopmental syndromes and developmental ciliopathy (Mascibroda *et al*., 2020; Oegema *et al*., 2017). Similarly, mutations in BRAT1 have been associated with neurodevelopmental and neurodegenerative disorders manifested with varying degrees of clinical severity (Balasundaram *et al*., 2021; Mundy *et al*., 2016; Nuovo *et al*., 2022; Scheffer *et al*., 2019; Srivastava & Naidu, 2016; Srivastava *et al*., 2016). These include rigidity and multi-focal seizure syndrome, lethal neonatal rigidity, and epilepsy of infancy with migrating focal seizures (Balasundaram *et al*., 2021; Mundy *et al*., 2016; Nuovo *et al*., 2022; Scheffer *et al*., 2019; Srivastava & Naidu, 2016; Srivastava *et al*., 2016). In order to further delineate the interaction between BRAT1 with INTS11/INTS9 we examined the effect of several patient-derived mutations of BRAT1 on its interaction with INTS11/INTS9 heterodimer (Fig. 5a). Human BRAT1 is a protein of 821 amino acids in length (protein id: NP_689956.2, gene id: NM_152743.4), displaying sequence conservation among vertebrates (Fig. 5b). We selected three of the reported neural disease-associated BRAT1 mutations including E522K (Girard *et al*, 2020), V62E (Mahjoub *et al*, 2019) and del_P309-Q310 (Valence *et al*, 2019) and mapped these mutations on the predicted three-dimensional model for human BRAT1 (Fig. 5c). Additionally, using published structure for INTS11/INTS9 heterodimer, we derived the BRAT1/INTS11/INTS9 trimeric structure (Fig. 5d). In this model, we assume INTS11 and INTS9 interact with each other at their C-terminal regions as previously described (Albrecht *et al*., 2018; Wu *et al*., 2017), while BRAT1 is predicted to fold as a circular helical structure around the stem of the structure holding INTS11/INTS9 heterodimer (Fig. 5d). This structural model suggests that BRAT1 and INTS4 interaction interfaces with INTS11/INTS9 heterodimer partially overlap and consequently form two mutually exclusive complexes of INTS11/INTS9/INTS4 or INTS11/INTS9/BRAT1 (Fig. 5e shows the multiple clashes between BRAT1 and INTS4 proteins when overlaying the INTS11/INTS9/BRAT1 complex into the experimental structure of the cleavage module). This result is consistent with our inability to detect BRAT1 in the affinity-purified eluate of Integrator complex isolated through Flag-INTS4 or Flag-INTS6 which are enriched with core-Integrator subunits (Sup. Fig. 5). To assess the effect of BRAT1 mutations in the association with INTS11/INTS9 heterodimer, we ectopically expressed the wild type and three disease-causing Flag-BRAT1 mutations (E522K, V62E and del_P309-Q310) in HEK293T cells. While the wild type and the two amino acids deletion (P309-Q310) of BRAT1 show normal association with INTS11/INTS9, the missense mutations either completely (E522K) or partially (V62E) disrupts the association between BRAT1 and INTS11/INTS9 heterodimer (Fig. 5f). These results connect the molecular defect, loss of BRAT1/INTS11/INTS9 interaction, with the phenotypic manifestation of disease phenotype by two disease-causing BRAT1 mutations described in patients with neurodevelopmental disorders (Girard *et al*., 2020; Mahjoub *et al*., 2019).

**Figure 5.**
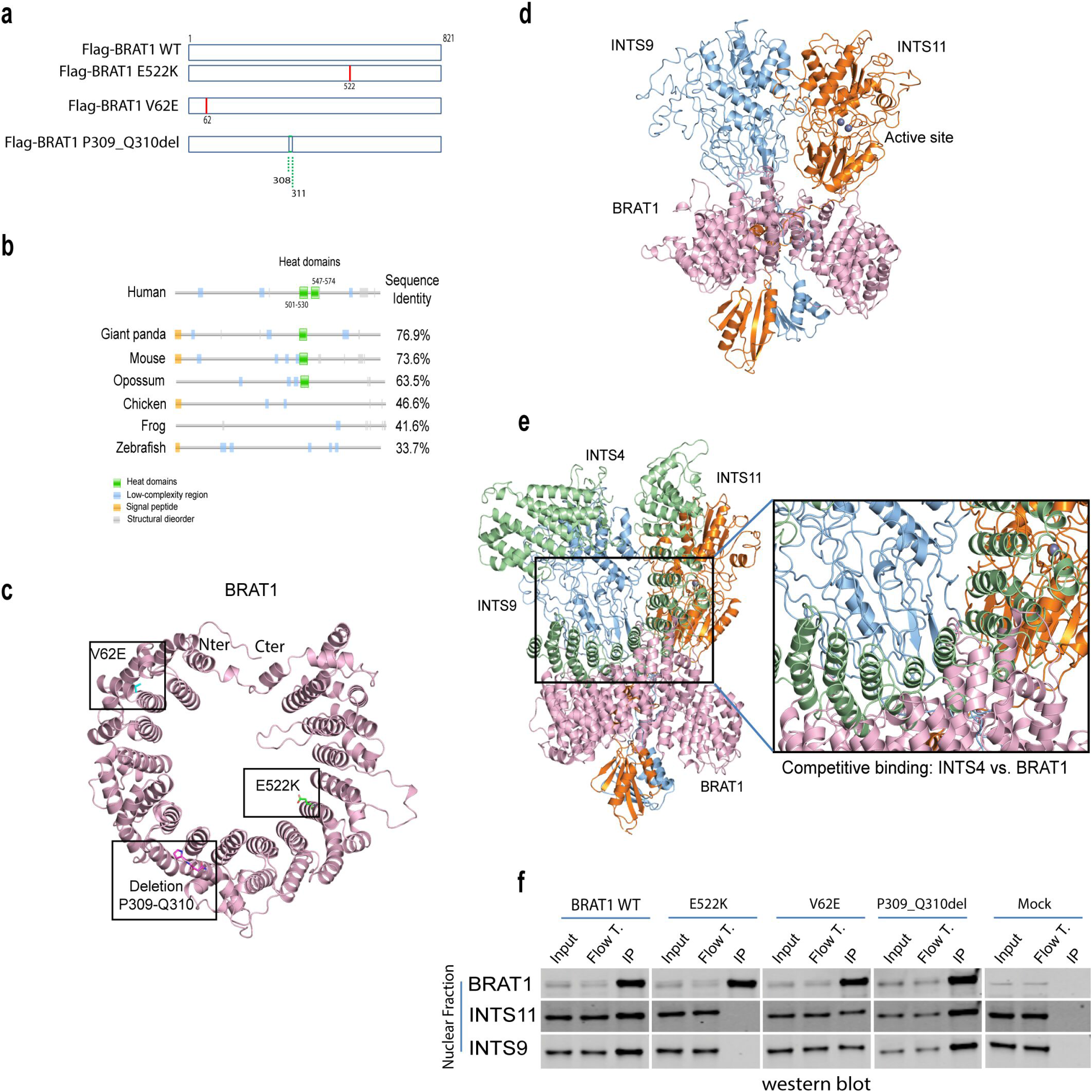
The pathogenic BRAT1 E522K mutation disrupts BRAT1 interaction with INTS11/INTS9 heterodimer. a, Schematic diagram showing the various constructs used for the investigation of interaction sites between BRAT1 and INTS11/INTS9 heterodimer. In addition to the wild-type BRAT1 (WT), three mutant constructs were engaged: E522K (c.1564G>A, p.Glu522Lys, red line), V62E (c.185T>A, p.Val62Glu, red line), and P309_Q310del (c.925_930del p.Pro309_Gln310del, green dash line). b, Protein sequence and domain conservation across representative vertebrate species. Pfam domains are shown as green boxes. Further properties are mapped to the sequence such as signal peptides in orange, low-complexity regions in blue and structural disorder in grey. Percentage of pairwise sequence identity to the human protein is shown on the right side. Pfam analysis shows that two HEAT domains are located at amino acids 501-530 and 547-574 ranges in human BRAT1. c, Tridimensional model of the human full-length BRAT1 protein (length=821 residues). The mutations under analysis were mapped to the structure and highlighted as sticks in different colors (green=E522, cyan=V62, pink=P309-P310del). N-terminal and C-terminal regions are labeled. d, Tridimensional model of the proposed binding mode between BRAT1 (pink), INTS9 (blue), and INTS11 (orange) obtained by molecular docking. The INTS11 active site contains two Zinc ions, shown as purple spheres. e, Overlay of the proposed tridimensional model of BRAT1/INTS9/INTS11 complex to the experimentally resolved cleavage model (INTS4/INTS9/INTS11)(Zheng *et al*., 2020) showing competitive binding between BRAT1 and INTS4. The detailed zoom on the right side illustrates the multiple clashes between BRAT1 and INTS4, rendering incompatible the interaction of BRAT1 with the full cleavage module and rest of the INTAC subunits. BRAT1, in pink, partially shares the binding interface of INTS4, in green, with the dimer INTS9-INTS11, in blue and orange respectively. f, Site directed mutagenesis (E522K, V62E, or P309_Q310del) was applied to BRAT1 protein by stable expression of Flag-BRAT1 mutants in HEK293T cells, and then its interaction with INTS11/INTS9 heterodimer was visualized by western blot. HEK293T cells stably expressing Flag-BRAT1 wild type (WT) isoform and mock were examined as positive and negative controls, respectively.

## Discussion

BRAT1 mutations in humans cause a spectrum of neurodevelopmental disorders often resulting in cerebellar atrophy and rigidity with or without seizures in neonates (Balasundaram *et al*., 2021; Mundy *et al*., 2016; Nuovo *et al*., 2022; Scheffer *et al*., 2019; Srivastava & Naidu, 2016; Srivastava *et al*., 2016). While BRAT1 protein is reported to associate with BRCA1 and ATM proteins (Aglipay *et al*., 2006), our biochemical isolation of BRAT1-containing complexes from human cells identifies INTS11 and INTS9 proteins as the only stably associated components of the BRAT1 complex (Fig. 1). This observation is consistent with a recent report describing BRAT1 interaction with INTS11/INTS9 heterodimer in human cells (Cihlarova *et al*, 2022). We find BRAT1 associates with INTS11/INTS9 heterodimer in both HEK293T (Fig. 1) and NT2 (Fig. 2a) human cell lines. Our biochemical analyses suggest that BRAT1 and INTS4 are components of distinct INTS11/INTS9-containing complexes. While INTS4 associate with INTS11/INTS9 to form the catalytic module of Integrator complex and allowing for the association of other core Integrator subunits (Zheng *et al*, 2020), BRAT1 binding to INTS11/INTS9 yields an exclusive trimeric complex. Our modeling of BRAT1 binding to INTS11/INTS9 supports this conclusion revealing a mutually exclusive binding interface between BRAT1 and INTS4 for binding the INTS11/INTS9 heterodimer (Fig. 5e). Importantly, the BRAT1 mutation of the key disease-causing residue E522K (Girard *et al*., 2020) present at the inner circular structure predicted to bind the C-terminal domain of INTS9/INTS11 completely abrogates BRAT1 interaction with INTS11/INTS9 highlighting the importance of this association in human cells.

A previous study suggests that mutation in BRAT1 causes the destabilization of its interaction with INTS11 and INTS9, thereby affecting the 3’-ends processing of UsnRNAs in U2OS cells (Cihlarova *et al*., 2022). They also found decreased levels of INTS11 in cell lines derived from patients with BRAT1 mutations. Therefore, using these cell lines, it would be difficult to assess whether BRAT1 directly contributes to RNA splicing defects seen following its depletion, since they may be caused by a secondary consequence of the INTS11 down regulation. Interestingly, in our case, the depletion of BRAT1 in NT2 cells does not affect the levels of INTS11 or INTS9, providing an opportunity to assess the functional consequence of BRAT1 perturbations in the absence of INTS11 loss. Our results suggest a specific effect of BRAT1 on the expression of a set of BRAT1-responsive neuronal genes distinct from the misprocessing of small nuclear RNAs by INTS11 depletion. Remarkably, while, we do not observe changes in INTS11 or INTS9 protein levels following loss of BRAT1 in NT2 cells, the depletion of BRAT1 in HeLa cells results in decreased levels of INTS11 protein (data not shown). Therefore, it is likely that BRAT1 contributes to INTS11/INTS9 protein stability in some cell lines.

Integrator is a multi-subunit protein complex implicated in multiple cellular pathways including processing of noncoding and coding transcripts as well as regulation of initiation and elongation of transcription (Beckedorff *et al*., 2020; Dasilva *et al*., 2021; Kim & Shiekhattar, 2016; Kirstein *et al*., 2023; Kirstein *et al*., 2021; Lai *et al*., 2015). Importantly, mutations in the number of Integrator subunits including INTS11 leads to brain developmental syndromes (Mascibroda et al., 2020; Oegema et al., 2017). These findings suggest an overarching role for Integrator as well as BRAT1/INTS11/INTS9 complex in the regulation of neuronal genes. Indeed, we find BRAT1 depletion results in the deregulation of a set of key neuronal identity genes during differentiation of NT2 cells. We show that BRAT1 plays an important role in recruiting INTS11 to these genes during their ATRA-mediated differentiation induction. Since many BRAT1-responsive genes are also targets of the neuronal silencer REST (RE1 Silencing Transcription Factor) (Duly *et al*, 2022; Sun *et al*, 2008), it is tempting to hypothesize that during neuronal differentiation BRAT1 plays a role in overcoming REST-mediated repression and therefore induce a critical set of neuronal identity genes. Taken together, our study provides a direct link between BRAT1 and INTS11/INTS9 heterodimer in regulation of neuronal genes during differentiation of human NT2 cells.

## Materials and methods

### Cell lines

HEK293T cell lines stably expressing Flag-BRAT1, Flag-INTS11, Flag-INTS6, and Flag-INTS4, and Flag-mock were established and cultured in DMEM containing puromycin (2.5 μg/ml) and supplemented with 10% FBS. NT2/D1 (NTERA-2cl.D1) cells, purchased from ATCC (CRL-1973), stably expressing Dox-inducible shControl and sh*BRAT1* were made by tet-pLKO-puro vectors purchased from Addgene (Cat# 21915). Dox (1 ug/ml) was freshly added in the NT2 culture media for BRAT1 depletion.

### Constructs

pLV puro EF1 HA-FLAG-BRAT11 transcript variant 2 (NM_152743.4) mutants including p.Glu522Lys (p.E522K, c.156G>A), p.Val62Glu (p.V62E, c.185T>A) and p.Pro309_Gln310del (p.P309_Q310del, c.925_930del) were synthetically made by VectorBuilder company (“https://en.vectorbuilder.com/”) and then the authenticity of the constructs were proved by sequencing.

### Affinity Purification of Flag-BRAT1, Flag-INTS11, Flag-INTS6, Flag-INTS4, and Flag-mock

HEK293 cell lines, stably expressing Flag-BRAT1, Flag-INTS11, Flag-INTS6, and Flag-INTS4 were cultured in DMEM media supplemented with puromycin, normocin and 10% FBS. Totally, sixty 15cm^2^ dishes of HEK293T cells (for each purification) harvested for performing the experiments. Following cytoplasmic fraction removal, nuclear lysate was extracted using 10 ml of the buffer contained 1.5 mM MgCl2, 20 mM Tris-HCl [pH 7.9], 0.5 mM DTT, 0.42M NaCl, 25% glycerol, 0.2mM EDTA, and 0.2 mM PMSF. The samples were dialyzed in 4L of BC150 buffer (20mM Tris pH 7.6, 0.2mM EDTA, 10% glycerol, 10mM 2-mercaptoethanol, 0.2mM PMSF and 150 mM KCl) for overnight at 4°C. Then, the complexes were purified from nuclear extracts using 1ml of anti-FLAG M2 affinity gel (Sigma #A2220) for overnight at 4°C. After washing twice with the 14 ml of BC500 buffer (20mM Tris pH 7.6, 10% glycerol, 0.2mM EDTA, 10mM 2-mercaptoethanol, 0.2mM PMSF and 0.5M KCl), and three times with the 14 ml of BC150 buffer for 10 minutes, the affinity columns were eluted with 500ul of 0.5ug/ul FLAG peptide prepared according to the manufacturer instruction (Sigma-Aldrich #F3290).

For size-exclusion chromatography, nuclear purified Flag-BRAT1 was loaded onto a Superose 6 (increase 10/300 GL) and equilibrated with BC300 buffer. The flow rate was kept at 0.35 ml/min. 0.5 ml fractions were collected and then TCA precipitated.

### TCA precipitation

90 ul of 100% TCA was added to 400ul of each fraction of HPLC fractionated samples and incubated for 15 min on ice. The samples were then centrifuged at 13,000 RPM for 30 min at 4°C and followed by washing twice with 800ul ice cold acetone. Pellet was dried on ice and resuspended in 50 ul of 1x Protein Sample Loading Buffer (LI-COR, #928-40004). 15 ul of the eluted samples were loaded on polyacrylamide gel and followed by western blot and silver staining.

### Western blotting

Cells were lysed by M-PER mammalian protein extraction reagent (Thermo Scientific, Cat# 78501), protein quantified by BCA protein assay ( Thermo scientific #23250), and 30 μg of the proteins were loaded on 4-12% Criterion TGX Stain-Free precast polyacrylamide gels (BIO-RAD, Cat# 5678084) and transferred to nitrocellulose membranes which were subsequently blocked by 5% BSA for 1 hour at RT. Membranes were incubated with primary antibodies (including: anti-INTS1 (Bethyl Laboratories, #A300-361A), anti-INTS3 (Sigma Prestige, #HPA074391), anti-INTS4 (Bethyl Laboratories, #A301-296A), anti-INTS6 (Novus, NB10086990), anti-INTS9 (Sigma Prestige, #HPA051615) anti-INTS11 (Sigma Prestige, #HPA029025), anti-INTS12 (Sigma Prestige, #HPA035772), anti-BRAT1 (Sigma Prestige, #HPA029455), anti-FLAG (Thermo Scientific, #MA1-91878-D680), anti-TUBB3 (Biolegend #801201), anti-GFAP (Invitrogen, #13-0300), anti-MAP2 (Millipore #AB5622), anti-Synapsin1 (Synaptic Systems #106 011C3) and anti-GAPDH (Abcam, #ab8245)) for overnight at 4°C. Then following 3 times washing for 10 min, blots were incubated with fluorescent-conjugated secondary antibodies for 1 hour at RT. Western blot results were visualized and quantitated by LI-COR ODYSSEY CLx system using ImageStudio software.

### Silver staining

Protein samples were loaded on a 4-20% Tris-glycine gel (Invitrogen), the gel was then fixed in 10% acetic acid and 50% methanol for 1 hour at room temperature. To complete fixation, the gel was transferred to 10% methanol and 7% acetic acid for 1 hour. 10% glutaraldehyde was used for washing the gel for 15 min followed by three times washing in MilliQ water for 15 min. Gel was stained for 15 min in 100ml of staining solution (1g AgNO3, 2.8ml NH4OH, 185ul NaOH (stock 10N in H2O), brought it up to 100 ml with MilliQ water). After 3 times washing for 2 min in MilliQ water, the gel was developed in 100 ml developing solution (52ul 37% formaldehyde and 0.5 ml 1% citric acid in 100 ml MilliQ water). 100 ml of a solution containing 50% methanol and 5% acetic acid was used to stop the reaction.

### Mass spectrometry

Following Flag-INTS11 and Flag-BRAT1 IPs in HEK293T cells, 50 ul of each final eluted samples were sent directly for sequencing. Samples were digested in-solution with trypsin and cleaned up using C18 spin column. Tryptic digests were analyzed using a standard 1.5 hr LC gradient on the Thermo Q Exactive HF or the Orbi Exactive HF mass spectrometers.

### Mass spectrometry analysis

MS data were searched with full tryptic specificity against the UniProt Human database(Bateman & Pilkington, 2011) using MaxQuant 1.6.17.0(Cox & Mann, 2008). MS data were also scanned for the common protein N-terminal acetylation, Met oxidation and Asn deamidation. Protein and peptide false discovery rates were set at 1%. Label-free quantification (LFQ) normalization was done using the MaxLFQ algorithm(Cox *et al*, 2014). We used Perseus (version 2.0.3.0(Stefka *et al*, 2016)) following the default protocol for label-free interaction data analysis. First, we filtered out reverse proteins, proteins identified only by one site, and contaminants. We log2 transformed the LFQ intensities and applied a protein filter requiring at least 2 non-zero values in one of the groups (IP vs. mock, 2 replicates each). We further used an imputation method to replace the missing values from a normal distribution (width factor=0.3 and down shift=1.8). Significant interactors were identified by two sample t-tests with s0=2 and permutation-based FDR<0.001 (number of randomizations=250).

### NT2 differentiation

NTERA-2 cl.D1 [NT2/D1] cells was purchased from ATCC and used for neural differentiation by a protocol as published previously by Peter W. Andrews(Andrews, 1984) with some modifications. Briefly, 5_X_10^6^ cells were seeded per 75 cm^2^ flask in Dulbecco’s Modified Eagle’s Medium (DMEM, Gibco, #11965-084) supplemented with 1X sodium pyruvate (Thermo Scientific #11360070), 10% fetal bovine serum (FBS, Atlas Biologicals, #F-0500-D), and normocin antibiotic (InvivoGen #ant-nr-2) at 37°C in 5% CO2. One day after seeding, cells were treated with 10^-5^ M all trans retinoic acid (ATRA, Sigma, Cat# R2625) for 42 days. Media was supplemented with fresh ATRA and exchanged every other day. Cells were then harvested at days 0 and 28 and used for RNA and protein extractions. For depletion of BRAT1, the cells were treated with 1ug/ml Dox (Selleckchem, #S4163) along with ATRA treatment.

### Immunostaining imaging

For immunostaining experiments, the proper number of the cells were cultured in 15 u-Slide 4 Well (ibidi# 80426). The cells were fixed in 4% PFA (4% PFA made in PBS) for 20 min at RT and followed by three times washing with PBS. The cells were then incubated with 1% Triton X/PBS for 20 min at RT and washed with PBS three times. Blocking was completed with 3% BSA protease free (Roche Ref# 03 117 332 001) for 1 hour at RT. 400 ul of each primary antibody diluted in 3% BSA protease free in PBS was added to each well and incubated overnight at 4°C and then washed with PBS three times. 400 ul of diluted secondary antibodies were added to each well and incubated for 30 min at room temperature and followed by three times washing with PBS. Finally, the cells were incubated with 8 ug/ml Hoechst in PBS and stained for 5 min and followed by three washes with PBS.

### RNA extraction, DNase treatment and cDNA synthesis

Total RNA was extracted using Trizol reagent according to the manufacturer’s protocol (Thermo Fisher Scientific, #15596026). RNA samples were treated by Turbo DNase RNase free Kit (Invitrogen, #AM1907) to remove DNA. 1ug of total RNA was used for cDNA synthesis by RevertAid First Strand cDNA Synthesis Kit (Thermo Scientific, #K1622) according to the manufacturer’s instructions.

### Reverse Transcriptase quantitative PCR (RT-qPCR)

1ul of cDNA (synthesized by cDNA synthesis kit, Thermo Scientific) was used for each RT-qPCR reaction with 1ul of 10 pmol of each primer (Table 1), 10 uL of SYBR Green Supermix (BioRAD) in a final volume of 20 ul, using a CFX96 real-time system (BioRAD). Thermal cycling parameters were: 3 minutes at 95°C, followed by 40 cycles of 10 s at 95°C, 20 s at 60°C followed by 30 s at 72°C. Raw data was analyzed by 2^-ΔΔCT^ method. Samples were run in triplicate. GAPDH was used as the internal control gene.

### Chromatin Immunoprecipitation quantitative PCR (ChIP-qPCR)

Prior to fixation, cells were washed twice with cold PBS, fixed with 1% fresh formaldehyde solution for 10 min rotating at room temperature and finally quenched by incubating with 125mM glycine rotating for 10 min at room temperature. Cells were washed twice with cold PBS, scraped and spined down at 2500 RPM. The cells were resuspended in 1000ul sonication buffer (20mM tris pH8, 50mM NaCl, 0.1% SDS, 0.5% Triton-x, 1mM EDTA) and rotated for 15 min at 4°C. Nuclei were isolated by spinning down 1 min at 5000 RPM. This step was repeated twice more, and the pellet then was resuspended in 1000ul sonication buffer and transferred to a Covaris 1ml millitube and sonicated in Covaris M-Series using peak power 75W, 10% Duty Factor, and 200 Cycles/Burst condition for 45 min. Samples were spined down at highest speed for 5 min at 4°C, and supernatant was transferred into low protein binding tube. Protein concentration was measured using Pierce BCA Protein Assay Kit (Thermo Scientific, #23250). 2 ug of antibodies was pre-bonded to 20ul of Protein A/G ChIP-grade Dynabeads in 100ul PBS/0.5% BSA overnight at 4°C. The beads were then washed twice with PBS/0.5% BSA. Almost 1-2mg of nuclei protein extract was incubated with pre-bound antibodies overnight at 4°C. ChIP was performed using antibodies for INTS11 (Sigma Prestige, #HPA029025), BRAT1 (Sigma Prestige, #HPA029455) and IgG (Rb IgG Isotype Control (Invitrogen, # 02-6102, LOT# R1238244, 5mg/ml). The IP samples were washed twice with Mixed Micelle Washed Buffer (150mM NaCl, 20mM Tris pH8, 5mM EDTA, 5.2% W/V Sucrose, 1% Triton-X, 0.2% SDS) for 10 min at 4°C, twice with Buffer 500 (0.1% sodium deoxycholate, 500mM NaCl, 50mM HEPES pH7.5, 1mM EDTA, 1% Triton-X) for 10 min at 4°C, twice with LiCl/Detergent (0.5% sodium Deoxycholate, 250mM LiCl, 1mM EDTA, 0.5% NP40, 10mM Tris pH8) for 10 min at 4°C and twice with 1x TE. The beads were resuspended in 75ul De-crosslinking buffer (20mM Tris pH8, 50mM NaCl, 0.5% SDS, 1mM EDTA) and incubated at 65°C 1200 RPM shaking for 30 min. Supernatant was transferred into a 1.5ml tube. De-cross linking was repeated twice more by adding 75 and 50 ul De-crosslinking buffer, and finally all 200 ul sample was de-crosslinked overnight at 65°C. The samples were then incubated with 1 mg/ml RNAse A (Invitrogen, #12091-021) and 1 mg/ml Proteinase K (Invitrogen, #AM2548) at 37°C for 40 min and at 65°C for 3 hours, respectively. After adding LiCl to 975mM, DNA was extracted by Phenol/Chloroform, and eluted in 40 ul of 1x TE buffer. The concentration of the samples was measured by Qubit. The same amount of DNA per sample was used for the ChIP-qPCR experiments.

### Co-Immunoprecipitation (Co-IP)

Culture medium was carefully removed from the cells and then cells were washed twice with ice cold PBS. 500µl ice cold IP lysis buffer (25 mM Tris-HCl pH 7.4, 150 mM NaCl, 1% NP-40, 1 mM EDTA, 5% glycerol add 50µl halt protease and phosphatase inhibitor TG270513 lot #1861282) was added to the cells and incubated on ice for 20 minutes with periodic mixing. The lysate was then transferred to a microcentrifuge tube and centrifuged at ∼13,000 g 10 min 4°C. Supernatant was transferred to a new tube. Protein concentration was measured by Pierce BCA Protein Assay Kit (Thermo Scientific, #23250). 15µl beads (protein A/G magnetic beads, Cat# 26126, Lot# TE261519; binding capacity 0.4µl/µl) per each IP was washed twice with 500µl Lysis buffer, resuspended in 100µl Lysis buffer with halt protease and phosphatase inhibitor and then coupled to antibodies at RT for 1h while rotating. 4 µg of INTS11 (Sigma Prestige HPA029025; lot#B117712), BRAT1 (Sigma Prestige, #HPA029455), INTS9 (Sigma Prestige, #HPA051615) or IgG (Rb IgG Isotype Control, Invitrogen, # 02-6102, LOT# R1238244, 5mg/ml) antibodies were used per IP. Bead coupling continued at 4°C for another hour. Pre-bound antibodies were washed once by lysis buffer. 2 mg of lysate was added to pre-bound antibody, rotating overnight at 4°C. Beads were washed twice with 500µl of lysis buffer. 42 µl of 1x Protein Sample Loading Buffer (LI-COR, #928-40004 diluted in applied lysis buffer) was added to the beads and incubated at 90°C 10min. 20 ul was loaded on 4-15% TGX gels (BIO-RAD).

### RNA sequencing (RNAseq)

500 ng of the total RNA extracted using TRIzol reagent (Thermo Fisher Scientific, #15596026) and treated with DNase RNase free Kit (Cat#AM1907) was used for the library preparation. The quality of total RNA was assessed by running it on 1% agarose gel and visualized by ethidium bromide staining. The RNA samples were depleted from ribosomal RNA and used for the library preparation using the TruSeq Stranded Total RNA Library Prep Kit (Illumina, #20020596) and then sequenced on Illumina NovaSeq to at least 45 million reads (single end reads, sequence length=100bp).

### RNA-seq analysis

Raw FASTQ data were processed with Trimmomatic(Bolger *et al*, 2014) (v. 0.38) removing adapters and low-quality reads. Reads were aligned to the human genome (hg19) using spliced transcripts alignment to a reference (STAR)(Langmead *et al*, 2009) aligner v. 2.5.3a with default parameters. RNA-seq by expectation-maximization (RSEM)(Li & Dewey, 2011) v1.2.28 was used to obtain the raw and normalized counts per gene against the human Ensembl reference transcriptome (release 87). Gene expression is normalized as Transcripts Per Million (TPM). Heat maps of the Z-score row-normalized gene expression and the unsupervised hierarchical clustering were generated using *clustermap* (method = *ward*, metric = *euclidean distance*)(Waskom). We determined the differential gene expression during neural differentiation in normal conditions and upon Brat1 depletion between day 0 and day 28 of the shRNA BRAT1 NT2 cells for either -Dox or +Dox condition using DESeq2(Love *et al*, 2014) at a 1.5-fold change cutoff (FDR<0.05). RNA-seq data (2 replicates per experimental condition) are available at Gene Expression Omnibus under accession number GSE237396. Enrichment analysis was performed on the differential expressed genes upon BRAT1 depletion by Enrichr(Chen *et al*, 2013; Kuleshov *et al*, 2016) against GO biological process.

### Protein conservation

Protein domain information was retrieved from Pfam database (version 35)(Mistry *et al*., 2021). Protein sequence identities were obtained by clustalo(Sievers *et al*, 2011) using Uniprot IDs of representative vertebrate species: Human=Q6PJG6, Giant panda=D2I4M3, Mouse=Q8C3R1, Opossum=F6TGD0, Chicken=E1BWY6, Frog=F6VIU7, and Zebrafish=Q1RLU1.

### 3D modeling

The tridimensional structure of the full-length, human BRAT1 protein (length=821 residues) was generated by Alphafold model prediction(Jumper *et al*, 2021; Varadi *et al*, 2022) and retrieved from https://alphafold.ebi.ac.uk with the accession code AF-Q6PJG6-F1. We mapped the human mutations under analysis to this structural model. Using HDOCK(Yan *et al*, 2020), we docked the BRAT1 model to the INTS9/INTS11 dimer obtained by Cryo-EM (PDB:7cun)(Zheng *et al*., 2020). We further compared the interaction interfaces of BRAT1 and INTS4 with the INTS9/INTS11 dimer via superimposition of the generated trimeric model into the cleavage module of the experimental structure by structural alignment of the common subunits with the open-source version of pymol (Schrödinger, L. & DeLano, W., 2020. PyMOL, Available at: http://www.pymol.org/pymol). The 3D figures were generated with the open-source version of pymol.

### Statistical analysis

Significant differences for ChIP-qPCR, RT-qPCR data, and average number of cell clusters microscopy analysis were determined by unpaired t-test using GraphPad Prism analysis (*p<0.05, **p<0.01, ***p<0.001).

## Acknowledgements

We thank the proteomics and metabolomics facility at Wistar Institute for sequencing of the MS samples, and Sylvester Comprehensive Cancer Center Onco-Genomics Core Facility for high-throughput sequencing of the RNA-seq samples. We thank current and past members of the Shiekhattar Lab for discussions and preliminary studies on BRAT1/Integrator functions. This work was supported by funding from University of Miami Miller School of Medicine, Sylvester Comprehensive Cancer Center and grants R01GM078455 from the National Institute of Health to R.S. Research reported in this publication was supported by the National Cancer Institute of the National Institutes of Health under Award Number P30CA240139. The content is solely the responsibility of the authors and does not necessarily represent the official views of the National Institutes of Health.

## Author contributions

S.D. performed RNA-seq, ChIP-qPCR, RT-qPCR, western blot, Flag affinity purifications, Imaging, stablishing sh*BRAT1* and shControl NT2 cells, neural differentiation of NT2 cells, endogenous co-immunoprecipitations, and silver staining experiments. H.G.D.S performed the bioinformatic analyses (RNA-seq, Mass spectrometry, and three-dimensional modeling). M.V. and S.D. performed Flag-BRAT1 mutants affinity purification and shControl and sh*BRAT1* NT2 cells establishment. H.A., 1. M. V. and S.D designed the mutant vectors. S.D. and R.S designed the experiments. S.D., H.G.D.S and R.S. wrote the manuscript.

## Disclosure and competing interest’s statement

The authors declare no competing interests.

## Data availability

All data needed to evaluate the conclusions in the paper are present in the paper and/or the Supplementary Materials. All raw and processed RNA-seq data are deposited in GEO under the accession number GSE237396.

## Code availability

All are available in the methods.

## Supplementary Information

is available for this paper

## Correspondence

should be addressed to R.S.

## Expanded figure’s legends

**Sup. Fig. 1. Integrator integrity is intact following BRAT1 depletion in NT2 cells.** Western blot of BRAT1 and Integrator complex subunits in NT2 cells 3 days after BRAT1 depletion shows that the integrity of Integrator complex is not affected by BRAT1 depletion. Cells are not ATRA-treated.

**Sup. Fig. 2. ATRA-mediated changes in gene expression upon normal differentiation of NT2 cells.** a, Volcano plot of ATRA-treated cells (without dox treatment) at day 28 compared to control NT2 cells at day 0, with down-regulated genes in blue, up-regulated genes in red and non-significant gene changes in black. 5687 (49%) genes were down-regulated, and 5883 (51%) genes were up-regulated (significance cutoffs: 1.5-fold change and FDR<0.05).

b, Gene enrichment of the up-regulated genes against Gene Ontology biological process 28 days after BRAT1 depletion in differentiated NT2 cells compared to cells expressing normal level of BRAT1. The GO analysis revealed that none of top ten gene ontology categories were related to brain development processes.

**Sup. Fig. 3. Other Integrator subunits such as INTS4 and INTS6 do not form a stable complex with BRAT1.** Western blot for nuclear purified Flag-INTS4 and Flag-INTS6 showing these subunits do not interact with BRAT1 in HEK293T cells.

